# Neutrophilic Inflammation in Models of Bronchopulmonary Dysplasia and Chronic Obstructive Pulmonary Disease is Rescued by a Lactobacilli Based Live Biotherapeutic

**DOI:** 10.1101/2023.03.02.529882

**Authors:** Teodora Nicola, Nancy Wenger, Xin Xu, Michael Evans, Gabriel Rezonzew, Luhua Qiao, Youfeng Yang, Namasivayam Ambalavanan, J Edwin Blalock, Amit Gaggar, Charitharth Vivek Lal

**Affiliations:** Division of Neonatology, Department of Pediatrics, University of Alabama at Birmingham, Birmingham, AL, United States; Division of Pulmonary, Allergy and Critical Care Medicine, Department of Medicine, University of Alabama at Birmingham, AL, United States

## Abstract

Bronchopulmonary dysplasia (BPD) is a chronic lung disease of prematurity. Exposure to noxious stimuli such as hyperoxia, volutrauma, and infection in infancy can have long-reaching impacts on lung health and predispose towards the development of conditions such as chronic obstructive pulmonary disease (COPD) in adulthood. BPD and COPD are both marked by lung tissue degradation, neutrophil influx, and decreased lung function. Both diseases also express a change in microbial signature dominated by *Proteobacteria* abundance and *Lactobacillus* scarcity. However, the relationship between pulmonary microbial dysbiosis and the mechanisms of downstream disease development has yet to be elucidated. We hypothesized that a double-hit hyperoxia and LPS murine model of BPD would show heightened Ac-PGP pathway and neutrophil activity. Through gain- and loss-of-function studies in the same model we showed that Ac-PGP plays a critical role in driving BPD development. We tested a novel inhaled live biotherapeutic using active *Lactobacillus* strains to counteract lung dysbiosis in *in vitro* and *in vivo* models of BPD and COPD. The *Lactobacillus* LBP is effective in improving lung structure and function, reducing neutrophil influx, and reducing a broad swath of pro-inflammatory markers in these models of chronic pulmonary disease. Live inhaled microbiome-based therapeutics show promise in addressing common pathways of disease progression that in the future can be targeted in a variety of chronic lung diseases.

---

The lungs are not sterile and instead harbor a low biomass microbiome (1). Bronchopulmonary dysplasia (BPD) is a chronic lung disease of premature infancy that results in an imbalance in lung tissue homeostasis and relatively immature repair systems within the developing lung (2-4). Chronic obstructive pulmonary disease (COPD) development can be linked to early life exposure to maternal cigarette smoke, respiratory illness, hyperoxia, and premature birth (5, 6). Neutrophilic inflammation marked by increased protease activity and driven in part by increased proteobacteria has been recognized as a primary contributor to the development of chronic lung diseases of infancy and adulthood (7-11). However, the mechanisms by which this inflammation can be reduced have not yet been determined in the context of disease treatment. Here we show that a live biotherapeutic product (LBP) containing a blend of commensal *Lactobacillus* strains reduces neutrophilic inflammation in models of chronic lung diseases such as BPD and COPD. We previously identified that infants with severe BPD had decreased firmicutes, specifically *Lactobacilli*, in their lungs with a corresponding increase in markers of neutrophilic inflammation (12). These findings were further tested in *in vitro* and gnotobiotic and humanized *in vivo* models of lung diseases (13, 14). This potent blend of live *Lactobacillus* strains was then formulated and engineered by spray drying for effective inhaled delivery into the distal regions of the lung. Here we demonstrate that inhaled *Lactobacillus* LBP is effective in reducing neutrophilic inflammation and improving lung structure through the MMP-PGP pathway in mouse models of BPD and COPD, performing comparably to inhaled steroid fluticasone furoate. Our results demonstrate the potential applicability of commensal *Lactobacilli*-based inhaled LBPs as a therapeutic approach to counter neutrophilic inflammation and tissue damage in chronic lung disease.

## Materials and Methods

### Human cohort study for the role of Ac-PGP in severe BPD

In a prospective cohort study, performed between October 2014 and December 2016, TA samples were collected at the UAB Regional Neonatal Intensive Care Unit from infants with established severe BPD at 36 weeks post menstrual age (PMA) as defined by the physiologic definitions were included (15). Tracheal aspirate samples were also obtained from gestational age-matched full-term infants, who were intubated at or within 6 hours of birth due to either surgical indication (congenital heart disease, abdominal wall defect) or perinatal depression (with no signs of meconium aspiration or other lung pathology). The samples were collected at the time of intubation prior to surgical procedures or anesthesia administration and served as controls. See supplement for collection and analysis details.

#### Study Approval

The protocol was approved by the IRB of the University of Alabama at Birmingham. A waiver of consent status was granted because the samples were obtained during routine clinical care and handled in a deidentified manner. All animal protocols were submitted and approved by the Institutional Animal Care and Use Committee of the University of Alabama at Birmingham. They were consistent with the Public Health Service policy on Humane Care and Use of Laboratory Animals (Office of Laboratory Animal Welfare, 2002).

### Animal Studies

C57BL/6 mice were obtained from The Jackson Laboratory (Bar Harbor, ME). All experiments were performed with a minimum of 6-8 mice to detect a 25% difference with 80% power based upon previous studies. See supplement for details on animal pulmonary function and sample processing.

#### Double hit BPD murine model

Newborn mice were randomized to treatment groups with PBS control or incremental intranasal doses of *Escherichia coli* 055: B5 LPS (Millipore Sigma). Pups were treated on postnatal day 3 (PN3) with 1mg/kg, postnatal day 6 (PN6) with 2.5mg/kg, postnatal day (PN9) with 5mg/kg, and postnatal day 12 (PN12) with 7.5mg/kg. Pups were also randomized to exposure to normoxia (21% FiO_2_) or hyperoxia (85% FiO_2_) from PN3-PN14. This represents the period of maximal alveolarization (16). Surrogate dams were switched every 24 hours from hyperoxia to normoxia to avoid oxygen toxicity. Daily animal maintenance was performed with exposure of the animals to room air for <10 minutes per day. A standard mouse pellet diet and water were provided ad libitum (16). Pups were euthanized for tissue harvest on PN14.

#### Murine postnatal Ac-PGP gain of function model

Newborn mice were randomized to treatment with normal saline control versus Ac-PGP (100µg/25µl saline). They received a total of 4 doses intranasally at PN3, PN6, PN9, and PN12. Pups were exposed to 21% FiO_2_ or 85% FiO_2_ from PN3-PN14 and were euthanized for tissue harvest at PN14.

#### Murine postnatal Ac-PGP loss of function model

Newborn mice that were exposed to 85% FiO_2_ and proteobacteria were randomized to treatment with arginine-threonine-arginine (RTR) versus normal saline at PN3, PN8, and PN13. The neutralizing Ac-PGP antagonist RTR, which functions to bind PGP both *in vitro* and *in vivo*, was synthesized by Anaspec and administered intraperitoneally. Daily animal maintenance was performed as described above.

#### Lactobacillus LBP treatment in double hit BPD model

Newborn pups were exposed to 85% FiO_2_ in addition to proteobacteria LPS (double hit model as described above) and randomized to treatment with *Lactobacillus* LBP or treatment with PBS. Mice were inoculated intranasally with the LBP (1×10^8^ total CFU) or PBS at PN3, PN6, PN9, and PN12 after 12 hours of LPS administration. Daily animal maintenance was performed as described previously.

#### In vivo strain survival and L (+) LA production

To determine whether administering live bacteria would be feasible from both dosing and viability perspectives, the lungs of 14 healthy 10–12-week-old female C57BI/6 mice were inoculated with 5×10^7^ cells/mL of the live bacterial blend (AB101, AB102, AB103). Two mice were sacrificed at each time point: immediately after inoculation, 2, 4, 8, 12, 24, and 72 hours after dosing. Lung tissue and bronchoalveolar lavage (BAL) fluid were sampled and assessed for their bacterial signature. Lung tissue with or without BAL was homogenized in 1 mL PBS and plated on agarose plates in serial dilutions of 1 µL, 10 µL, and 100 µL. BAL was mixed with 500 µL PBS and plated on agarose plates in serial dilutions of 0.2 µL, 2 µL, and 20 µL. Plates were incubated for 16 hours at 37ºC overnight and colonies were counted visually. L (+) lactic acid production in the lung tissue and BAL was also measured using a Lactate Colorimetric Assay (MAK064, Sigma).

#### Cigarette smoke COPD model

20 6 to 8-week-old female C57BL/6 mice were exposed to cigarette smoke or air and treated with a blend of AB101, AB102, and AB103 strains or saline control. Mice who received smoke were exposed for 3 hours a day Monday through Friday over a 1-month period (see supplement). The blend of strains was administered at a concentration of 1×10^8^ CFU suspended in 50 μL of PBS. Mice were anesthetized and dosed IT with the *Lactobacillus* blend or 50 μL saline (control) every other day on Monday, Wednesday, and Friday (MWF). Mice were sacrificed after 4 weeks.

#### Porcine pancreatic elastase COPD model

6 to 8-week-old C57BL/6 mice were allocated into 7 groups (6 females, 6 males per group). The animals were administered 0.25 IU porcine pancreatic elastase (PPE) on Day 1 and Day 11 via oropharyngeal instillation to induce emphysema development over the course of the 3-week experiment. As injury, mice either received 50 µL PBS as a negative control, PPE, or a double-hit exposure of PPE and 1 hour later, 100 µg of *E. coli*-derived lipopolysaccharide (LPS). LPS administration to PPE mice models severe airway dysbiosis as seen in exacerbations of COPD.

#### Lactobacillus LBP co-treatment of PPE model mice

Beginning on Day 2, mice were treated with 1.5 mg of the *Lactobacillus* LBP containing 1×10^8^ CFU live bacteria via Insufflator three times a week on Monday, Wednesday, and Friday (MWF) for 3 weeks post-induction. The 1.5 mg absolute dose was chosen based on the results of a pilot dose tolerability test. Control groups with PBS, PPE, and PPE+LPS received 50 µL of saline dosed every other day in lieu of the LBP. One group was given LBP daily to test if a more rigorous dosing frequency was tolerable by the mice compared to the MWF schedule. All animals were sacrificed 21 days after first PPE induction.

#### Lactobacillus LBP post-injury treatment of PPE model mice

Following the design of the PPE model described above, mice began therapeutic treatment starting on Day 12 only after PPE injury had been established. Mice were treated MWF with either PBS (control), LBP, or the inhaled corticosteroid fluticasone furoate (Sigma). Animals were sacrificed 21 days after first PPE induction.

#### Statistical analysis

All data were expressed as mean ± SEM. One-way ANOVA was used to analyze the effect of LPS, LPS + hyperoxia, and LBP combination on animals. Similarly, multiple-comparison testing (Student-Newman-Keuls) was performed if statistically significant (\**P*<0.05) was noted by ANOVA. GraphPad Prism version 7.00 for Mac OS X (GraphPad Software, La Jolla, California, USA, www.graphpad.com) was used for all data analysis.

## Results

### Airways of premature infants with severe BPD are marked by increased *Proteobacteria* and decreased Lactobacilli, increased neutrophilic inflammation, and increased levels of matrikines

Our group recently performed a prospective cohort study in the Regional Neonatal Intensive Care Unit at UAB during which tracheal aspirates were collected from infants with a diagnosis of severe BPD as well as gestational age-matched full term control infants (mean postmenstrual age of 38.3±2.2 weeks). We discovered infants with severe BPD have increased proteobacteria abundance and proteobacteria endotoxin concentrations in their airways (Figure 1A) (12). In order to further understand the potential link between dysbiosis and tissue destruction, markers of neutrophil activity including myeloperoxidase (MPO) and neutrophil elastase (NE) were analyzed. The tracheal aspirates from patients with severe BPD showed higher MPO and NE concentrations when compared to controls, indicating a significant neutrophil influx (Figure 1B-C). In addition, MMP-9, prolyl endopeptidase (PE), and Ac-PGP were increased in tracheal aspirates from patients with severe BPD as compared to controls (Figure 1D-F).

**Figure 1.**
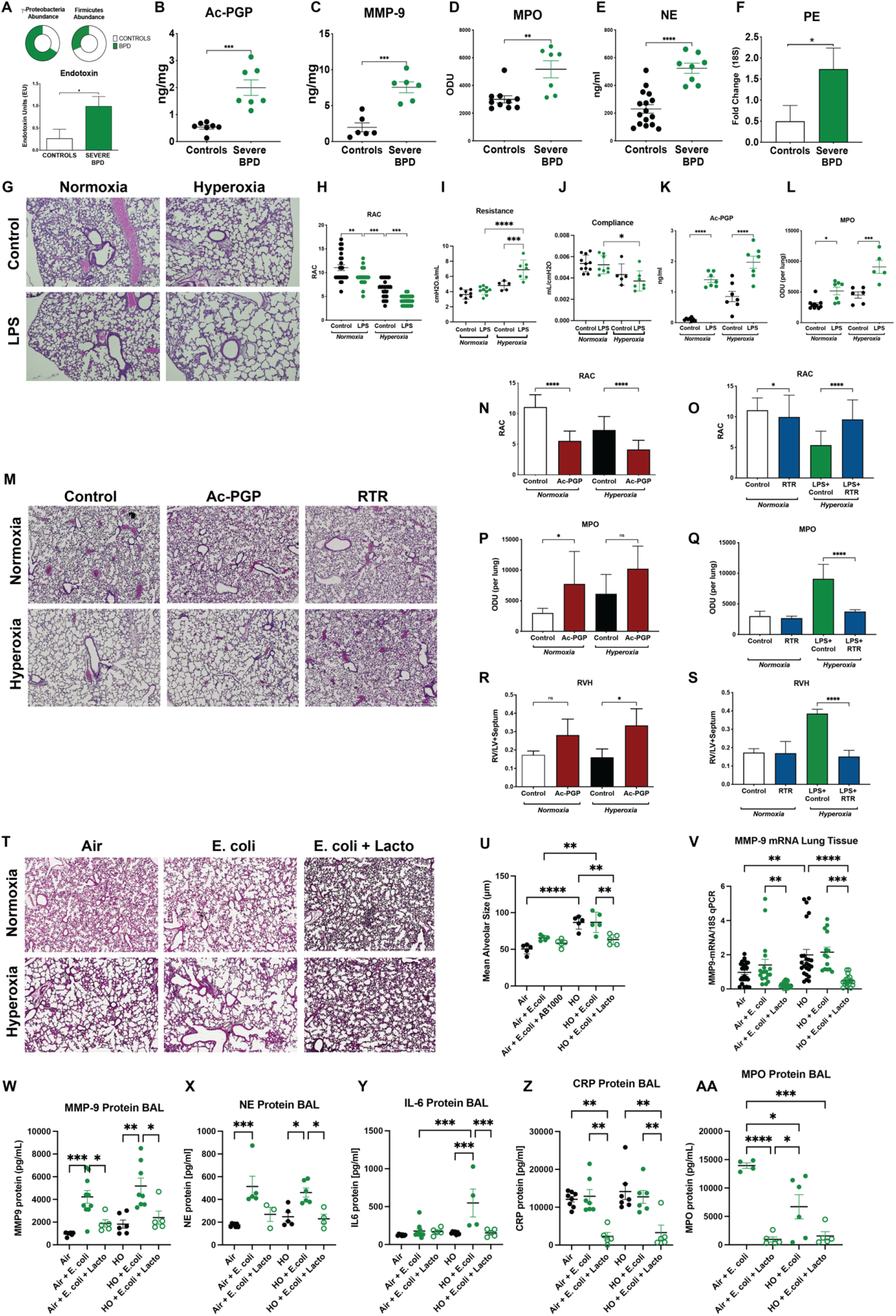
Severe BPD is marked by increased proteobacteria and neutrophil influx mediated by the MMP-9/Ac-PGP pathway. **<u>Human</u>** **A)** Infants with severe BPD had increased *Proteobacteria*, decreased *Firmicutes*, and increased endotoxin levels compared to control infants (*P<0.05). Infants with severe BPD had significantly higher concentrations of **B)** MPO, **C)** NE, **D)** MMP-9, **E)** prolyl endopeptidase (PE), and **F)** Ac-PGP in their tracheal aspirates than control infants. **<u>Hyperoxia + LPS Mice (Double-Hit)</u>** **G)** Mice exposed to hyperoxia in combination with LPS resulted in severe alveolar hypoplasia and simplification. Tissue slices from harvested lung tissue were photographed at a magnification of 100X after staining with hematoxylin-eosin. **H)** Radial alveolar count (RAC) was significantly reduced in LPS-exposed mice when exposed to normoxia or hyperoxia. Pulmonary function worsened as measured by **I)** resistance (increased) and **J)** compliance (decreased) upon exposure to hyperoxia and LPS. In both normoxia and hyperoxia, LPS-exposed mice had significantly higher concentrations of **K)** Ac-PGP protein and **L)** myeloperoxidase (MPO) protein in the bronchoalveolar lavage fluid (BAL). **<u>Hyperoxia + Ac-PGP/RTR Mice</u>** **M)** Ac-PGP and LPS exposure each in combination with hyperoxia resulted in severe alveolar hypoplasia and simplification. Treatment with RTR (arginine-threonine-arginine), an Ac-PGP inhibitor, resulted in improved alveolar structure. 4x magnification. **N)** RAC was significantly reduced upon dosing with Ac-PGP or LPS and **O)** significantly improved upon treatment with RTR in normoxia and hyperoxia. **P)** MPO expression was significantly increased upon Ac-PGP exposure in normoxia. **Q)** MPO expression was significantly increased upon exposure to LPS + hyperoxia and significantly reduced upon RTR treatment. **R)** Right ventricular hypertrophy (RVH) was significantly increased in Ac-PGP mice exposed to hyperoxia and **S)** significantly reduced in LPS + hyperoxia mice treated with RTR. ***<u>Lactobacillus</u>* <u>LBP Treatment of Double-Hit Mice</u>** **T)** Hyperoxia exacerbated alveolar simplification in mice exposed to *E. coli* intranasally while intratracheal administration of live biotherapeutic improved tissue structure in *E. coli* + hyperoxia mice. 4x magnification. **U)** Mean alveolar size was increased upon hyperoxia and *E. coli* exposure and restored to normal levels upon treatment with LBP. **V)** MMP-9 mRNA expression in lung tissue significantly increased upon *E. coli* exposure and further increased in hyperoxia. LBP reduced expression in normoxia and hyperoxia. **W)** MMP-9 protein in the BAL significantly increased upon exposure to *E. coli* and decreased upon treatment with LBP in both normoxia and hyperoxia. **X)** NE protein in the BAL significantly increased upon exposure to *E. coli* and decreased upon treatment with LBP in hyperoxia. **Y)** Hyperoxia and *E. coli* exposure significantly increased IL-6 protein in the BAL and was decreased by LBP treatment. **Z)** CRP protein in the BAL was significantly decreased by LBP treatment in normoxia and hyperoxia. **AA)** MPO protein in the BAL was significantly decreased by LBP treatment in normoxia. Bars represent mean + SD. \**P*<0.05, \*\**P*<0.01, \*\*\**P*<0.001

### Double hit mice model of severe BPD shows significant alveolar hypoplasia and increased matrikine surge

As the tracheal aspirates from infants with severe BPD exhibited both proteobacteria abundance and neutrophil influx with increased chemoattractant elements, we sought to further define the role of airway dysbiosis in the pathogenesis of BPD. In order to accomplish this end, we created a double-hit murine model of BPD by intranasally instilling LPS into C57BL/6 mouse pups with the added insult of hyperoxia exposure from PN3-PN14. As observed in the histologic images in Figure 1G, the mouse pups exposed to both LPS and hyperoxia exhibit the most severe alveolar hypoplasia when compared with control animals in hyperoxic and normoxic conditions. On assessment of radial alveolar counts (RAC), the mouse pups exposed to both LPS and hyperoxia exposure showed significantly decreased alveoli (\**P*<0.05) when compared to control mice (Figure 1H). LPS alone also led to significantly lower alveolar counts when compared to control mice in normoxia (Figure 1H). Pulmonary function worsened upon exposure to hyperoxia and LPS as measured by increased resistance and decreased compliance (Figure 1I, J).

When assessing the concentration of Ac-PGP, LPS-exposed animals showed an increase when compared to PBS-treated animals in normoxia (Figure 1K). LPS-exposed animals and controls also showed higher concentrations of Ac-PGP when exposed to hyperoxia. MPO levels followed the same pattern, rising with LPS exposure alone and combined with hyperoxia (Figure 1L).

### Gain of function of matrikine Ac-PGP produces severe BPD, loss of function protects in mice

In order to further elucidate the role of Ac-PGP in the pathogenesis of BPD, newborn mice were dosed with exogenous Ac-PGP and exposed to normoxia or hyperoxia. The lung tissue was then examined to determine if these mice would exhibit the phenotype of severe BPD (Figure 1M). When radial alveolar counts were analyzed, it was discovered that mice treated with Ac-PGP had significantly reduced alveolar counts when compared to controls (Figure 1N). In addition, both control and Ac-PGP exposed mice showed decreased RAC when exposed to hyperoxia; however, the Ac-PGP exposed mice in hyperoxia showed even further reduced alveolar counts when compared to controls in hyperoxia.

Additionally, MPO was increased in mice exposed to Ac-PGP in both normoxia and hyperoxia when compared to controls (Figure 1P)). The control mice in hyperoxia showed increased MPO when compared to the control mice in normoxia, but the Ac-PGP exposed mice exhibited a significantly higher concentration of MPO in hyperoxia.

In order to assess the effect of Ac-PGP on pulmonary vascularization, echocardiograms were performed on the studied animals. In both the normoxia and hyperoxia exposure groups, treatment with Ac-PGP resulted in significantly more severe right ventricular hypertrophy (RVH) when compared with control animals (Figure 1R). These findings suggest that Ac-PGP can result in neutrophil influx and the development of a BPD phenotype.

We then analyzed the loss of Ac-PGP function with the hypothesis that loss of Ac-PGP function would be protective against the development of a BPD phenotype. In this case, mice were treated with an Ac-PGP antagonist known as arginine-threonine-arginine (RTR) in addition to LPS and hyperoxia exposure. We noted that the hyperoxia-exposed mice treated with LPS and RTR had significantly higher alveolar counts when compared to hyperoxia exposed mice treated with LPS alone (Figure 1O). MPO concentration was analyzed in order to gauge neutrophil activity after treatment with RTR. In hyperoxia, mice treated with LPS and RTR showed lower concentrations of MPO than mice exposed to LPS alone (Figure 1Q). Upon analysis of pulmonary vasculature, it was noted that hyperoxia exposed mice treated with LPS and RTR showed less severe RVH (Figure 1S). These findings suggest that treatment with an Ac-PGP antagonist may be protective, as it results in decreased neutrophil activation as well as reduced tissue damage.

### *Lactobacillus*-based LBP improves lung morphology and function in double-hit murine BPD model

Since our previous study revealed that infants with severe BPD have an increase in airway proteobacteria and a decrease in airway *Lactobacillus* abundance, we investigated treatment of an *in vivo* model of proteobacteria abundance and hyperoxia with the LBP blend. Using the double-hit BPD injury model as described above, we treated mice with the *Lactobacillus*-based LBP.

Mouse pups were dosed intratracheally with the LBP blend in order to determine if treatment directly to the lungs would counteract the lung injury typically observed in this model. Lung morphology was analyzed and revealed that the lung injury model had higher alveolar counts when treated with LBP, which suggests less alveolar hypoplasia and simplification (Figure 1T-U). When mice exposed to hyperoxia and *E. coli* were treated with LBP, the concentration of MMP-9 in lung tissue and MMP-9 and NE protein in BAL decreased, indicating a lower neutrophilic influx (Figure 1V-X). IL-6, CRP and MPO protein levels also decreased in hyperoxia + *E. coli* mice dosed with LBP (Figure 1Y-AA).

### Live *Lactobacillus* strain blend reduces MMP-9 levels more than individual strains

Given the key results linking lung tissue degradation and the state of the lung microbiome, we investigated opportunities to utilize commensal bacteria to reduce inflammation and damage. We exposed human bronchial epithelial cells (HBEC) to *E. coli* and/or hyperoxia to generate *in vitro* models of lung damage. Cells were treated with three live *Lactobacillus* strains individually and as blends of differing concentrations. When cells were treated with a blend of three *Lactobacillus* strains AB101, AB102, and AB103, IL-1β and MMP-9 levels decreased more than when treated with individual strains (Figure 2A-B).

**Figure 2.**
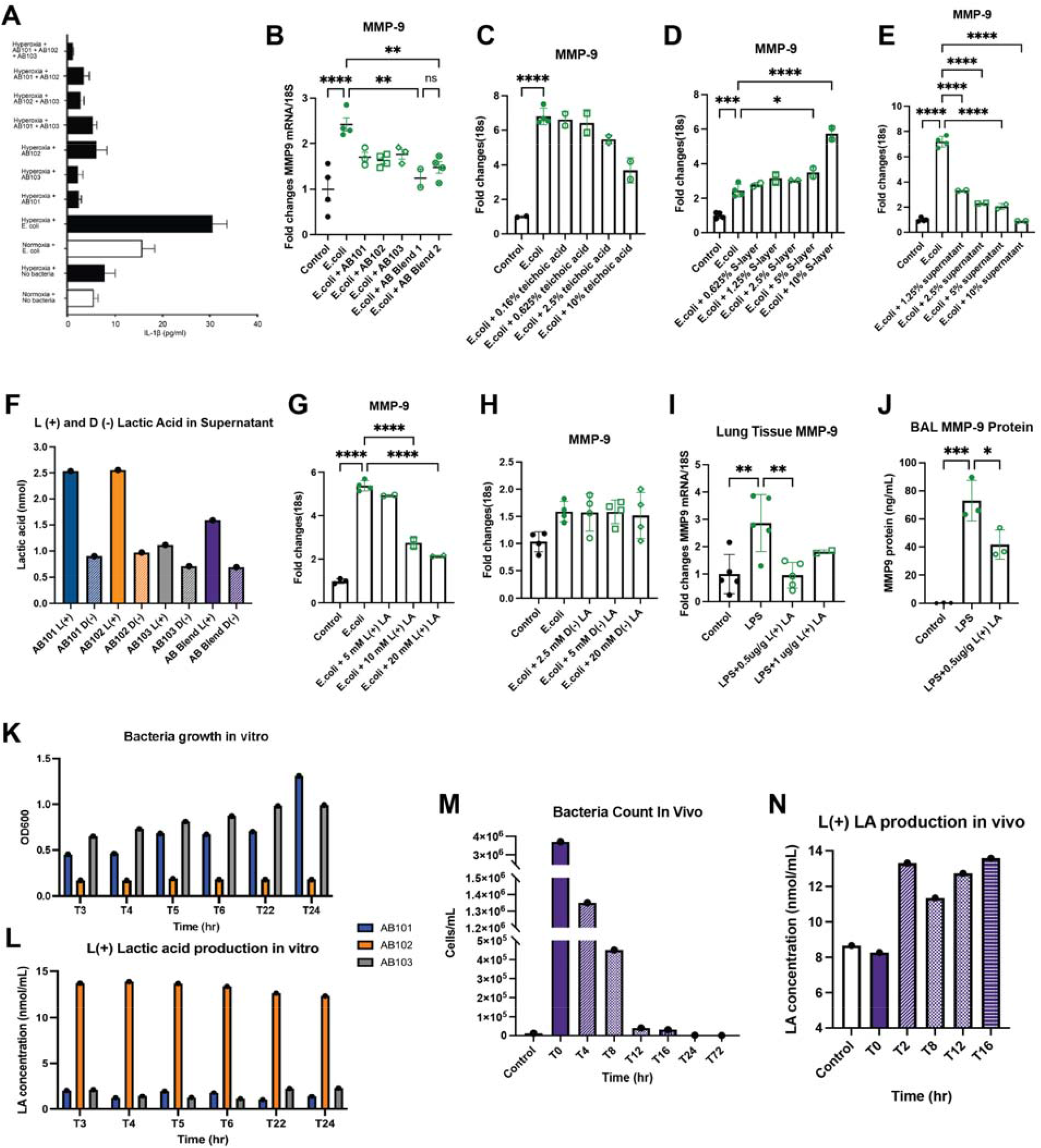
*Lactobacillus* blend reduces neutrophilic inflammation through L (+) lactic acid production. Live *Lactobacillus* strains AB101, AB102, and AB103 were assessed for their anti-inflammatory activity individually and as blends. The blend of all three strains performed better than individually in reducing **A)** IL-1β protein and **B)** MMP-9 protein in human bronchial epithelial (HBE) *in vitro* models of noxious stimuli exposure. *Lactobacillus* components **C)** teichoic acid and **D)** S-layer isolated from strains have no effect on reducing MMP-9 expression in HBE cells exposed to *E. coli*. **E)** *Lactobacillus* blend culture supernatant in increasing concentrations reduces MMP-9 expression. **F)** *Lactobacillus* strains produce more L (+) lactic acid than D (+) lactic acid individually and as a blend. **G)** L (+) lactic acid reduces MMP-9 expression in HBE cells exposed to *E. coli* while **H)** D (-) lactic acid does not. Mice exposed to LPS intratracheally who were dosed with L (+) lactic acid showed decreases in **I)** lung tissue MMP-9 expression and **J)** BAL fluid MMP-9 protein levels. Cell culture performed in triplicate. Bars represent mean + SD. \**P*<0.05, \*\**P*<0.01, \*\*\**P*<0.001, \*\*\*\**P*<0.0001

### Live bacteria active components reduce MMP-9

We then isolated cellular components and byproducts of the *Lactobacillus* strains to identify the potential anti-inflammatory agents at work. Isolated teichoic acid (Figure 2C) and peptidoglycan (Figure 2D) did not have any effect on MMP-9 expression in the *in vitro E. coli* HBEC model, while polysaccharide and S-layer (Supplemental Figure 1) in fact increased MMP-9 expression. Cell supernatant was effective in reducing MMP-9 expression in a dose-dependent manner (Figure 2E). *Lactobacilli* are known lactic acid producers, and we measured the L (+) and D (-) lactic acid output of each strain (Figure 2F). L (+) lactic acid was effective at reducing MMP-9 expression *in vitro* while D (-) was not (Figure 2G, H).

L (+) lactic acid dosed intratracheally to WT mice exposed to intratracheal LPS reduced MMP-9 expression in lung tissue (Figure 2I) and MMP-9 protein levels in BAL (Figure 2J) in comparison to mice who received LPS exposure without treatment.

### *In vivo* LBP lactic acid production and clearance

We first grew each *Lactobacillus* strain *in vitro* and measured the L (+) lactic acid output of each strain. While AB102 maintained a relatively low concentration (Figure 2K), it was the highest producer of L (+) lactic acid out of the three strains (Figure 2L). We then inoculated healthy WT mice intratracheally with a blend of the three strains to model inhalation and clearance of an LBP. While the cell count in the lungs steadily decreased over 72 hours as measured by colony counts of BAL plus lung tissue (Figure 2M), L (+) lactic acid production stayed relatively consistent through 16 hours (Figure 2N).

### Engineered LBP particles show favorable delivery and viability characteristics

Using the three live *Lactobacillus* strains, we engineered an inhalable particle that contains live bacteria spray dried with stabilizers and excipients with favorable characteristics for deep lung delivery. Several rounds of spray drying optimization (Table 1) yielded a lead formulation (1) as shown in scanning electron microscope images (Figure 3). As measured by gravimetric impaction, the formulation had a very small median mass aerodynamic diameter (MMAD) of 2.4 µm (Table 1). 74% of total powder mass was <3.3 µm, meaning that approximately 74% of the particles were small enough to be delivered to the distal lung. Viability of the bacteria in the particles was tested, and 99% was retained comparing bacteria counts pre and post processing. These tests validated that a live bacteria blend could be successfully engineered with favorable particle characteristics for deep lung delivery while maintaining excellent viability levels.

**Table 1.** Aerosol and physical characteristics of three spray dried powder formulations. MMAD – median mass aerodynamic diameter; GSD – geometric size distribution; FPF – fine particle fraction; gPSD – geometric particle size; DSC – differential scanning colorimetry; Tg – glass transition temperature; KF – Karl Fischer; CFU – colony forming units

|  | **1 – lead** | 1 – repeat run | 2 | 3 |
| --- | --- | --- | --- | --- |
| Delivered dose (mg) | **26.82** | 25.6 | -- | 26.32 |
| Delivered dose (% of 30 mg) | **88%** | 85% | -- | 86% |
| MMAD (µm) | **2.4** | 2.3 | -- | 2.5 |
| GSD | **1.7** | -- | -- | 1.6 |
| FPF<5.0µm | **94%** | 95% | -- | 94% |
| FPF<3.3µm | **73%** | 76% | -- | 74% |
| gPSD, 4 bar (µm)<br>D10/D50/D90 | **0.487/1.40/3.02** | -- | 0.602/1.35/2.74 | 0.459/1.38/3.00 |
| DSC Tg (Tg1/Tg2) | **40.8C/124.7C** | -- | 43.1C/121.3C | 39.6C/No data |
| Bulk density (g/cc) | **0.26** | -- | 0.28 | 0.19 |
| Water activity | **0.082** | -- | 0.079 | 0.094 |
| Moisture (KF) | **1.48%** | 1.48% | 1.38% | 1.46% |
| Viability (CFU/g) | **3.8x10<sup>9</sup>** | 4.6x10<sup>9</sup> | -- | 1.7x10<sup>9</sup> |

**Figure 3.**
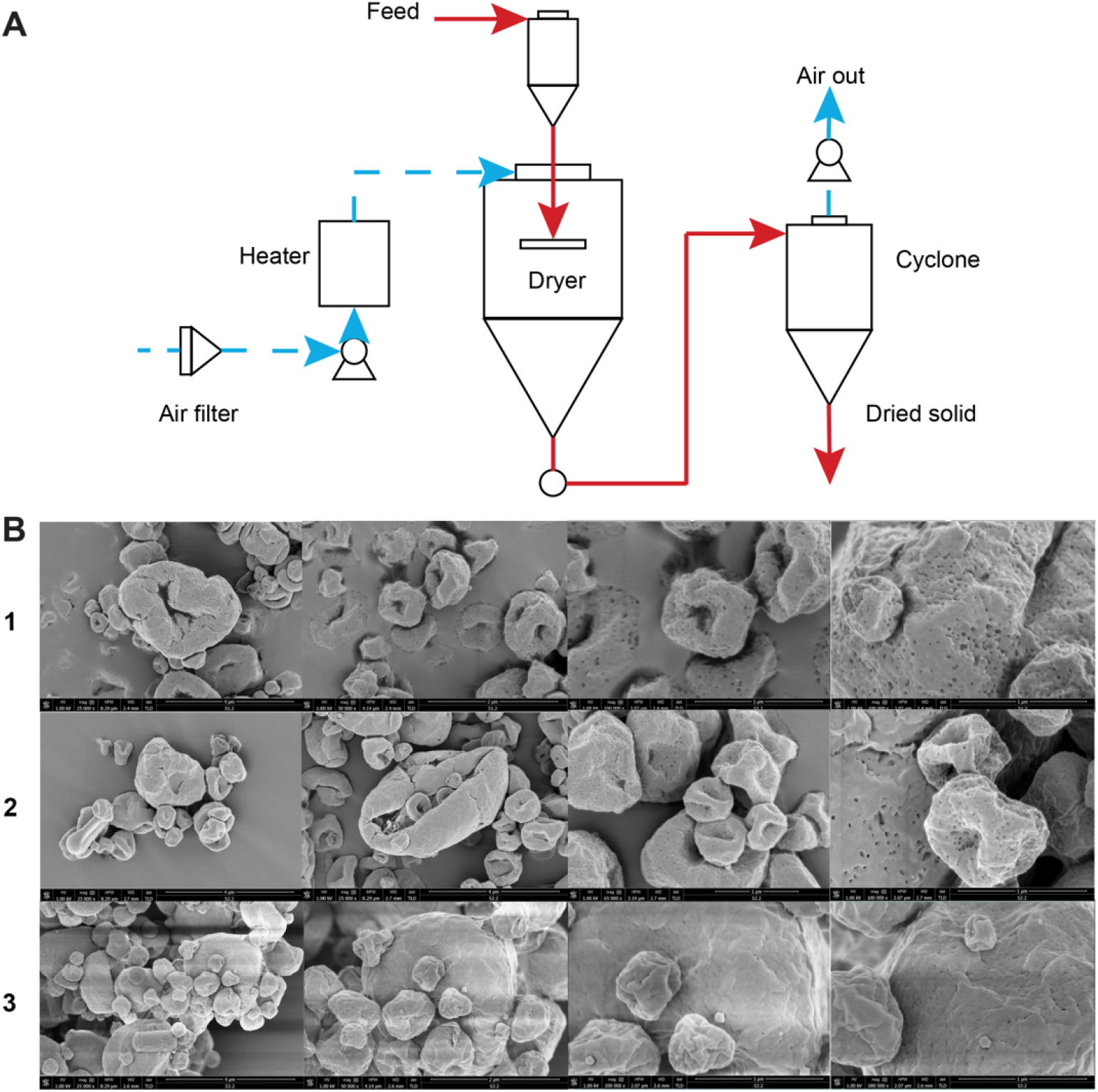
Spray drying produces small, flowable powder particles containing viable bacteria. **A)** Schematic of spray drying process used to dry *Lactobacillus* strains into small, flowable powder particles. **B)** Scanning electron microscope (SEM) images of three formulations of powder particles after drying. Images at 25,000, 50,000, and 100,000X magnification (left to right).

### Inhaled *Lactobacillus* LBP attenuates lung tissue damage and inflammation in murine models of COPD

Having generated a flowable powder for inhalation containing viable bacteria with anti-inflammatory action, we proceeded to test the *Lactobacillus* LBP in preclinical animal models of chronic lung disease. Based on the promising results with the LBP in the BPD double hit mouse model as well as mechanistic commonalities, we evaluated anti-inflammatory activity of the LBP in three mouse models of COPD.

First was an etiological COPD model using cigarette smoke. After 4 weeks of smoking, mice showed increased MMP-9 mRNA in lung tissue and protein in serum (Figure 3A-B). When treated with LBP, mice showed a drop in MMP-9 levels as well as an increase in protective IgA protein in the BAL compared to controls (Figure 4A-C).

**Figure 4.**
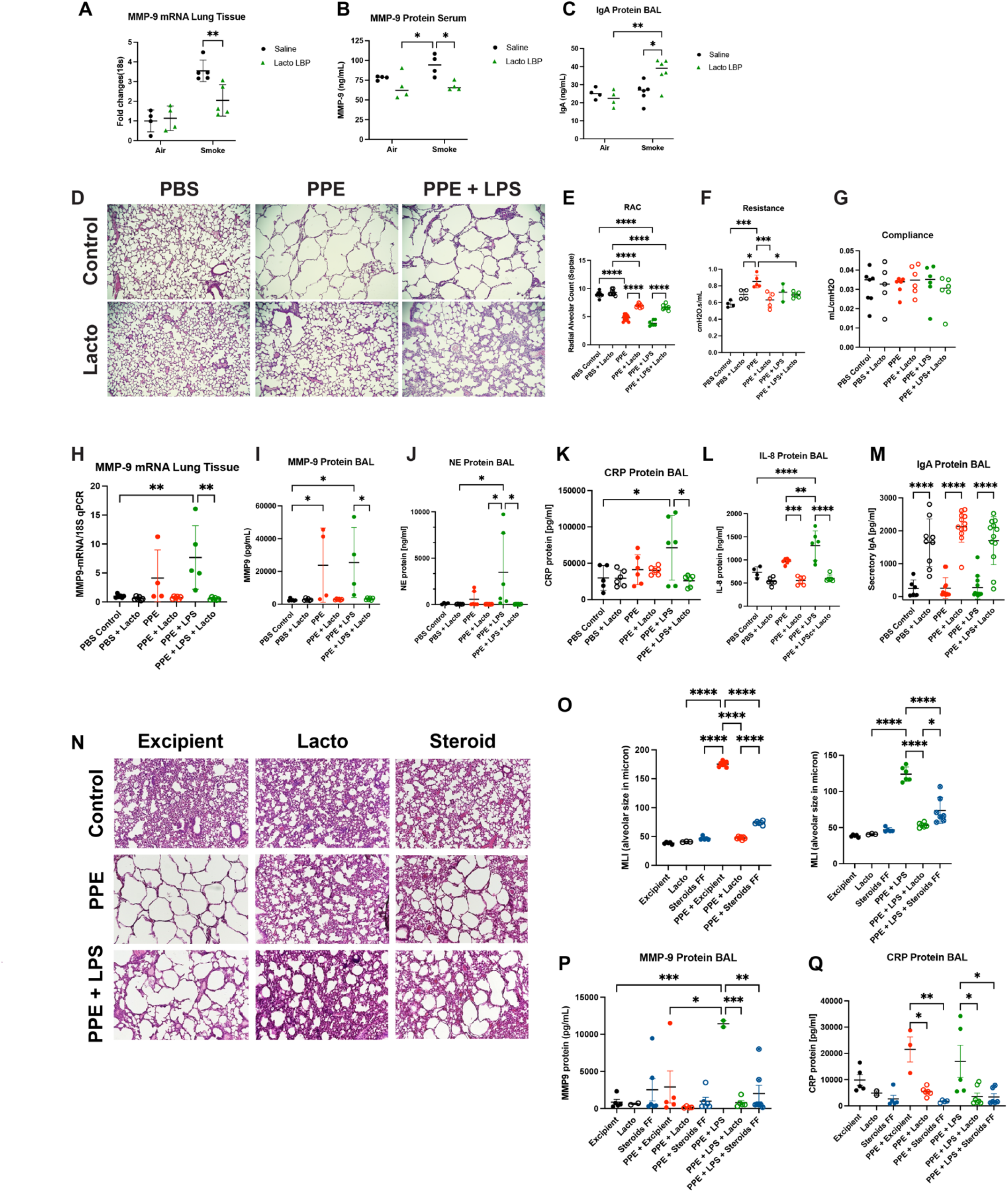
*Lactobacillus*-based LBP improves biomarkers of inflammation and lung structure and function in three murine models of COPD. In a cigarette smoke exposure model of lung injury, mice treated with a *Lactobacillus* LBP (Lacto) intratracheally showed improvements in **A)** lung tissue MMP-9 expression, **B)** serum MMP-9 protein levels, and **C)** BAL IgA protein levels. **D)** Intratracheal porcine pancreatic elastase (PPE) and LPS exposure leads to significant alveolar hypoplasia and simplification while inhalation of LBP concurrent with injury period reduces tissue damage. **E)** RAC was significantly improved upon treatment with *Lacto* LBP in PPE and PPE + LPS exposure groups. **F)** Lung function worsened as measured by resistance (increased) upon exposure to PPE and improved upon treatment with *Lacto* LBP. **G)** Lung function as measured by compliance showed no significant changes among groups. **H)** MMP-9 expression in lung tissue significantly increased upon PPE + LPS exposure and decreased upon treatment with *Lacto* LBP. **I)** MMP-9 protein in the BAL significantly increased upon PPE and PPE + LPS exposure and decreased upon treatment with *Lacto* LBP in PPE + LPS mice. **J)** NE protein in the BAL significantly decreased upon *Lacto* LBP treatment in PPE + LPS mice. **K)** CRP protein in the BAL significantly increased upon PPE + LPS exposure and decreased upon treatment with *Lacto* LBP. **L)** IL-8 protein in the BAL significantly decreased upon *Lacto* LBP treatment in PPE and PPE + LPS mice. **M)** IgA protein in the BAL significantly increased (improved) upon *Lacto* LBP treatment in control, PPE, and PPE + LPS mice. **N)** Intratracheal PPE and LPS leads to alveolar hypoplasia and simplification. *Lacto* LBP treatment after two weeks of injury is established reduces inflammation and improves tissue structure. Steroid treatment after injury decreases inflammation. **O)** PPE exposure (LEFT) and PPE + LPS exposure (RIGHT) each increases mean linear intercept and *Lacto* LBP treatment significantly decreases (improves) it. The steroid fluticasone furoate (FF) also significantly reduces MLI but not as much as *Lacto* LBP. **P)** *Lacto* LBP works better than the steroid in reducing MMP-9 protein in BAL of mice exposed to PPE + LPS. **Q)** *Lacto* LBP works as well as the steroid in reducing CRP protein in BAL of mice exposed to PPE + LPS. Bars represent mean + SD. \**P*<0.05, \*\**P*<0.01, \*\*\**P*<0.001, \*\*\*\**P*<0.0001

Next, we tested structural and structural + dysbiosis models. We first conducted a dose tolerability test to assess how much LBP powder PPE mice could tolerate. A single administration of 5 mg of the LBP directly to the lungs by insufflation was greater than the maximum tolerated dose, resulting in the reduction of the dose to 2.5 mg for the rest of the week. 5 mg of powder caused airway clogging and death based on lung volume limitations of the mice; the death was not associated with the bacteria (Supplemental Table 1). A dose of 2.5 mg was tolerated by the mice. To ensure the animals tolerated a duration of dosing longer than a week, a lower dose of 1.5 mg of the LBP administered every other day was selected for future studies.

In parallel 3-week experiments, mice were exposed to either PPE or PPE+LPS and treated concurrently with the LBP. Mouse lungs were fixed and sectioned for histology and imaged at 10x magnification (Figure 4D). Mice exposed to PPE showed a strong emphysema phenotype compared to controls. Mice exposed to both PPE and LPS showed the emphysema phenotype with additional inflammation in the lung tissue. Treatment with the LBP rescued lung structure compared to both injury groups. Increases in radial alveolar counts (RAC) in the LBP-treated groups confirmed the improvements observed visually (Figure 4E). Lung function worsened as measured by increased resistance upon exposure to PPE and improved upon treatment with the LBP (Figure 4F). There was no significant change in compliance (Figure 4G).

Lung tissue MMP-9 expression showed a significant increase upon exposure to PPE+LPS and significant decrease upon treatment with the LBP (Figure 4H). Mice exposed to PPE alone did not show a significant increase in MMP-9 transcription or decrease in response to LBP treatment.

MMP-9, NE, IL-8, C-reactive protein (CRP), and immunoglobulin A (IgA) were evaluated in BAL and serum. Treatment with the LBP resulted in significantly lower MMP-9, NE, CRP, and IL-8 protein in PPE+LPS dosing groups, and lower MMP-9 and IL-8 in the PPE groups (Figure 4I-L). Protective protein IgA levels in BAL were significantly increased in every group treated with the LBP (Figure 4M). Treatment with the LBP resulted in significantly lower MMP-9, NE, and CRP protein in serum (Supplemental Figure 2). Dosing the LBP every other day consistently produced lower levels of pro-inflammatory proteins.

### *Lactobacillus* LBP performs as well as steroid in post-injury treatment model of COPD

After showing improvement in inflammatory biomarkers and tissue structure with LBP treatment concurrent to injury, we treated mice after the injury was established. Mice were exposed to PPE or PPE+LPS and treated with either the LBP or the steroid fluticasone furoate to the lungs. Even when treating mice after PPE or PPE+LPS injury was established, LBP treatment resulted in significant improvements to lung tissue structure as measured by MLI (Figure 4N-O). Steroid treatment resulted in a modest improvement in structure consistent with typical mouse healing rates in a 3-week PPE model. The LBP exhibited impressive steroid-like anti-neutrophilic inflammatory action and worked better than and as well as, respectively, the steroid in reducing MMP-9 and CRP protein in BAL (Figure 4P, Q). In fact, the LBP performed better than steroids in reducing a variety of additional pro-inflammatory proteins in the BAL including MIP-3b, IL-11, IL-16, and TIMP1, among others (Supplemental Figure 3).

### *Lactobacillus* LBP shows favorable safety and biodistribution profile in PPE mouse model

To characterize the *Lacto* LBP’s inhaled safety profile, we tested the drug product in a respiratory safety and biodistribution study using PPE model mice. Mice were exposed to PPE or a PBS control as previously described and treated daily for 14 days with either nothing, placebo powder, 1 mg LBP, or 3 mg LBP by insufflator. On Day 21, half the mice had their vitals taken and were sacrificed in a terminal sacrifice. On Day 28, the second half of the mice had their vitals taken and were sacrificed in a recovery sacrifice. Vital measurements included body temperature, body weight, pulse distension, breath distension, heart rate, breath rate, and oxygen saturation. Across all groups in terminal and recovery, vital signs did not indicate a negative response to the inhalation of the LBP powder (Figure 5). Despite some significant changes between groups, the values did not exceed clinically normal ranges.

**Figure 5.**
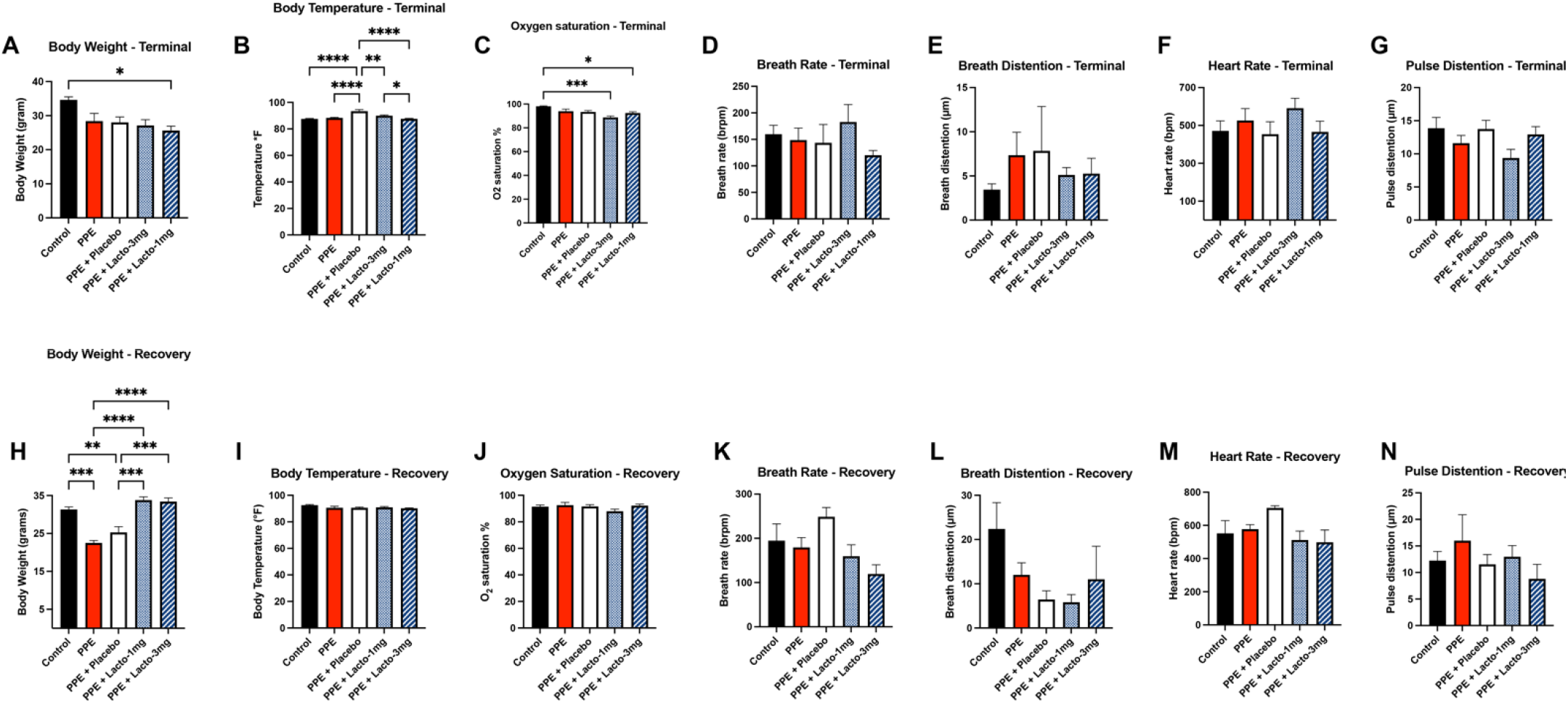
Inhaled *Lactobacillus*-based LBP shows favorable safety and biodistribution profile in PPE mouse model. Body weight, temperature, oxygen saturation, breath rate, breath distension, heart rate, and pulse distension remained in normal ranges after 2 weeks LBP dosing in **A-G)** terminal sacrifice mice and **H-N)** recovery sacrifice mice.

At terminal and recovery sacrifices, lung, brain, heart, liver, kidneys, spleen, sinuses, and serum were harvested. Each tissue had bacterial gDNA extracted and PCR run using primers specific to each *Lactobacillus* strain in the LBP (AB101, AB102, and AB103) to see if any API distributed beyond the target tissue of the lungs. PCR gels confirmed that no *Lacto* LBP remained in the lungs at either terminal or recovery sac, nor did it travel to distal tissues.

### Identifying additional pathways of interest affected by *Lactobacillus* LBP

To investigate further what pathways are at play upon LBP treatment in multiple injury models, we conducted a PCR array. HBE cells were exposed to *Pseudomonas*, cigarette smoke, smoke and *Pseudomonas*, or *E. coli* and treated with the LBP or a PBS control. Using custom primers for 88 human lung epithelial cell-specific genes, a PCR array was run, and a gene expression heat map generated (Supplemental Figure 3A). Of those 88 genes, 9 genes were significantly downregulated upon treatment with the LBP: TGFβ-1, CD36, CLDN18, GRAMD2, HSPA4, ICAM1, MMP-11, MMP-9, SCNN1B. MMP-9 served as the benchmark for the accuracy of our experiment. To crosscheck this result *in vivo*, we measured expression levels of these genes in the lung tissue of mice from the PPE+LPS concurrent treatment experiment. Of these 9 genes, all except TGFβ-1 and ICAM1 were significantly reduced by the LBP when treating the double injury PPE+LPS mice (Supplemental Figure 4B).

We additionally conducted Qiagen Ingenuity Analysis to assess the effect of LBP treatment in *in vitro* models of lung injury. *E. coli* and *Pseudomonas* appeared to use similar pathways with a seven gene overlap while cigarette smoke had only three genes in common with *E. coli* (Supplementary Table 2, Supplemental Figure 5).

## Discussion

While multiple studies have explored the role of Ac-PGP, ours is the first to our knowledge to determine the link between airway proteobacteria abundance and neutrophilic inflammation in the pathogenesis of BPD and COPD. In our *in vitro* experiments, exposure to *E. coli* resulted in elevated markers of neutrophilic inflammation. Similarly, LPS exposure *in vivo* resulted in elevated markers of neutrophilic inflammation as well as evidence of reduced alveolarization, which is consistent with the presentation of BPD (2). This was seen independent of hyperoxia exposure, although hyperoxia appeared to potentiate the effects. This is consistent with previous human studies showing that infants with severe BPD have an airway microbial signature that is defined by proteobacteria abundance and *Lactobacillus* deficiency (3, 12, 17).

The generation of Ac-PGP appears to be imperative to the development of BPD and COPD. This discovery was made through the generation of Ac-PGP gain and loss of function murine models. Through intranasal instillation of Ac-PGP, markers of neutrophilic inflammation (MPO) increased independently of hyperoxia. Worsening alveolar simplification (decreased RAC) and abnormal vascular development (increased RVH) were also noted. When treated with RTR (arginine-threonine-arginine), the markers of neutrophilic inflammation decreased. The murine models also showed an increase in the number of alveoli and a decrease in RVH, suggesting improved alveolar and vascular development. RTR binds and neutralizes PGP sequences and has been noted to attenuate lung damage in murine models of acute lung injury/acute respiratory distress syndrome as well as in allergic airway disease (18, 19).

Based on the importance of the presence of *Lactobacillus* in the lung microenvironment, we examined *Lactobacillus* strains for their anti-inflammatory activity. The live strains performed better in synergy rather than as individuals in reducing MMP-9 and IL-1β in *in vitro* models. IL-1β is a pro-inflammatory cytokine that has been found in increasing concentrations in the airway secretions of patients with BPD (20-23). We found that bacterial supernatant isolated from the blend of *Lactobacillus* strains was effective in reducing MMP-9 to a similar degree as live bacteria. *Lactobacilli* produce L (+) lactic acid which when isolated also reduced MMP-9 *in vitro*. Exosomal microRNA is another source of anti-inflammatory action that would be present in bacterial supernatant (24). When inhaled, live bacteria act as an efficient delivery mechanism for the multiple anti-inflammatory moieties that they produce. This is one of the benefits to using bacteria rather than the isolated components as the drug product.

In order to explore the efficacy of a *Lactobacillus*-based therapeutic, we treated the double hit murine model with the respiratory LBP. Following treatment with the live biotherapeutic, partial protection of lung structure and markers of neutrophilic inflammation (MPO, MMP-9, NE) was noted. These results suggest that supplementing commensal strains to the lung microenvironment may attenuate neutrophilic inflammation and reduce tissue damage associated with the development of BPD.

We then tested the LBP in multiple murine models of COPD. First was an etiological model of COPD using cigarette smoke exposure, second was a structural model using porcine pancreatic elastase (PPE), and finally a double-hit structural and dysbiosis model (PPE+LPS). Cigarette smoke exposure is a common noxious stimulant leading to COPD pathogenesis (25). PPE generates an emphysema phenotype in three weeks, making it an ideal preliminary short-term model to test the LBP’s efficacy *in vivo*. In addition, PPE+LPS models a more severe phenotype COPD with inflammation caused by proteobacteria abundance in the airways.

Improvements in both pro and anti-inflammatory biomarkers in the BAL of LBP-treated COPD mice across models suggest that inhalation of the *Lactobacillus*-based blend successfully reduced inflammation in the lung microenvironment whether treated concurrently or post injury. The commensal *Lactobacillus* in the formulation exerted an anti-inflammatory effect that decreased known pro-inflammatory markers MMP-9, NE, CRP, and IL-8 while elevating anti-inflammatory marker IgA. Similar trends were seen in the serum as well, suggesting that the anti-inflammatory action of the LBP in the lungs can trigger signaling cascades on a systemic level. Our PCR array and subsequent pathway analysis also strongly indicate that the LBP has a broad therapeutic effect not limited to the Ac-PGP/MMP-9 cascade. Potential biomarker targets showed reduced levels upon LBP treatment including CD36 (thrombospondin-1 receptor), CLDN18 (claudin 18), GRAMD2 (GRAM domain containing 2), HSPA4 (heat shock 70 kDa protein 4), MMP-11, and SCNN1B (sodium channel, non-voltage gated 1 beta subunit, aka βENaC). CD36 is a receptor to thrombospondin (TSP-1) which is involved in converting the anti-inflammatory cytokine TGFβ-1 to its active form (26). CLDN18 is a tight junction protein involved in maintaining proper alveolar barrier function (27, 28), and GRAMD2 is enriched in alveolar epithelial cells (AT1) (29). SCNN1B is involved in mucus clearance, and overexpression may result in mucus stasis, inflammation, and increased risk of infection (30, 31). Common themes among these targets are ECM regulation and lung microenvironment homeostasis.

Perhaps the most interesting finding was the favorable performance of the LBP compared to fluticasone furoate, an FDA-approved inhaled corticosteroid used in the COPD triple therapy Trelegy. The LBP reduced MMP-9 and other pro-inflammatory cytokines as well as, and in some cases better than, the steroid in the PPE + LPS COPD model. Prolonged use of steroids for COPD exacerbations is associated with side effects including increased risk of pneumonia, skin thinning, sepsis, and death (32, 33). The safety and biodistribution study in the PPE emphysema model indicated that inhalation of the LBP did not initiate adverse reactions in diseased mice. Furthermore, the bacteria strains did not translocate to distal tissues or accumulate in the lungs, supporting the hypothesis that the bacteria have a localized yet transient presence in the airways. In the future, live biotherapeutics may reduce or replace the need for inhaled steroids by virtue of their strong anti-inflammatory action combined with a stronger safety profile and additional potential structural benefits.

In conclusion, the pathogenesis of chronic lung diseases is complex and multifactorial in nature. Based on previous studies, proteobacteria abundance and the Ac-PGP axis play heavy roles in development of BPD and COPD. Neutrophilic influx creates a pro-inflammatory milieu in the lungs which results in tissue damage. Inhaled *Lactobacillus*-based live biotherapeutic shows promising therapeutic potential as evidenced by efficacy in our robust set of BPD and COPD models.

While symptomatic treatments are available for patients, there remains a lack of preventative or curative therapies for BPD or COPD. Performance comparable to inhaled steroid has massive implications in affecting the landscape of standard of care for COPD patients by providing a novel supplemental or replacement treatment option. These and future studies provide foundational research that may lead to improved care for patients with these chronic and devastating diseases.

## Author Contributions

Conceptualization: CL, NA, AG, NW, TN, JB. Experimentation: TN, XX, ME, GR, LQ, YY. Analysis and interpretation: NW, TN, ME, LQ, YY, CL. Drafting and editing manuscript: NW, TN, CL, NA. Funding acquisition and research supervision: CL, AG.

