## Supplemental Methods and Figures for "Neutrophilic Inflammation in Models of Bronchopulmonary Dysplasia and Chronic Obstructive Pulmonary Disease is Rescued by a Lactobacilli Based Live Biotherapeutic"

**Human Tracheal Aspirate**

*Human Tracheal Aspirate Collection:* Tracheal aspirate samples were obtained from established BPD patients and gestational age-matched full-term controls as described above. The samples were obtained after ensuring adequate oxygenation. The protocol for obtaining tracheal aspirates involved instilling 1mL of sterile isotonic saline into the infant’s endotracheal tube, manual bagging through the endotracheal tube for 3 breaths, and suctioning of the fluid into a sterile mucus trap. The sample was centrifuged at 3,000 *g* for 10 minutes in order to separate the supernatant and cell pellet. They were frozen at -80°C until further processing.

*Analysis of Ac-PGP by electrospray ionization-liquid chromatography-tandem mass spectrometry:* Acetylated PGP was purchased from Bachem (Torrance, CA) and purified to neutrophil chemotactic activity. The acetylated PGP in tracheal aspirates was measured using a MDS Sciex API-4000 spectrometer (Applied Biosystems, Carlsbad, CA) equipped with a Shimadzu HPLC (Columbia, MD). HPLC was performed suing a 2.0_150-mm Jupiter 4u Proteo column (Phenomenex, Torrance, CA) with Buffer A (0.1% HCOOH) and Buffer B (MeCN+0.1% HCOOH). HPLC was initially performed with 95% Buffer A and 5% Buffer B from 0-0.5 minutes, then increased over 0.5-2.5 minutes to 0% Buffer A/100% Buffer B. Background was removed by flushing with 100% isopropanol/0.1% formic acid. Positive electrospray mass transitions were at 312-140 and 312-112 of acetylated PGP (1, 2).

***Lactobacillus* LBP Formulation**

The *Lactobacillus* LBP material was developed in collaboration with iPharma Ltd (San Francisco, CA). The blend contains *Lactobacillus* strains AB101, AB102, and AB103 spray dried with proprietary excipients and stabilizers. *Lactobacillus* strains AB101, AB102, and AB103 are licensed from the American Type Culture Collection (ATCC).

**Cell Culture and Treatment:**

Human bronchial epithelial cells (HBEC) were used (American Type Culture Collection, PCS-300-010). Each experiment was performed in triplicate.

*Hyperoxia model:* Cells were exposed to 21% FiO_2_ to represent normoxic conditions or 85% FiO_2_ in a hyperoxia cell culture chamber to represent hyperoxic conditions. Cells were exposed to 1.0μg/ml of *Escherichia coli* 055: B5 LPS (Millipore Sigma) or PBS as control.

*E. coli in vitro model:* 6-well plates were inoculated with proteobacteria (*E. coli*), the *Lactobacillus* blend, or individual *Lactobacillus* strains AB101, AB102, and AB103 or isolates. *E. coli* (*E. coli* serotype K1) was grown on Luria Bertani (LB) broth containing 100μg/ml of rifampin (Millipore Sigma). 1x10^6^ CFU of each bacteria strain AB101, AB102, and AB103 were inoculated in 100μl of antibiotic and serum free media. Media without bacterial inoculation was used as a control. The supernatant (200μl) was collected at 12 hours from each well for each experiment.

**Animal Studies**

*Cigarette smoke administration in COPD model:* Mice who received smoke were exposed for 3 hours a day Monday through Friday over a 1-month period. To smoke the mice, cigarette smoke from reference cigarettes RF2F (Tobacco and Health Research Institute, University of Kentucky) were diluted 1:1 and introduced into a chamber. Smoke was generated using the SIREQ generator provided with the inExposure system. The nicotine concentration in the smoke using the above parameters was approximately 11μg/l across all mice.

*LBP powder tolerated dose test:* In preparation for dosing mice with representative LBP drug product test material in a porcine pancreatic elastase (PPE) model of emphysema in COPD, several dose volumes of the *Lacto* LBP and dosing frequencies in mice with administration by Insufflator were tested. The Insufflator device is the representative equivalent of a DPI used in these mice. The representative test material was a neat formulation containing spray dried research grade bacteria with 2.0% w/w polysorbate 80 with no other excipients. This batch of material contained 2.4x10^10^ cells/mL of the *Lactobacillus* blend. 27 female C57BI/6 mice were used in this 5-day experiment. On Day 1, mice were inoculated intranasally with either 0.25 IU of PPE, 100 µg LPS, or 50 µL PBS (control). 1 hour later, mice were Insufflated with 5 mg of LBP containing 1x10 8 CFU of live bacteria. Three groups of mice served as control and did not receive any doses of representative LBP. PPE and LPS only groups were included as positive controls to verify that any deaths were not caused by the severity of the injury induced by either.

Mice who received the Day 1 administration of 5 mg dose of LBP struggled due to its high physical volume rather than the quantity of bacteria contained within. The powder clogged their airways and 1-2 mice in both the PPE and control groups died (Groups 1, 2, 4). To account for the poor survival rate, the dose was lowered to 2.5 mg (0.5x10^8^ CFU) for the rest of the dosing period.

Mice receiving *Lacto* LBP dosing continued through Day 8; half received LBP daily and half received LBP every other day. Mice were dosed with another 0.25 IU of PPE, 100 µg of LPS, or 50 µL of PBS on Day 8, 1 week after the first exposure. On Day 9 following the last LBP administration, all surviving mice were sacrificed, and lung tissue was collected for quantitation of MMP-9 mRNA levels via qPCR.

*Safety and biodistribution in PPE emphysema model:*

We conducted a respiratory safety and biodistribution study in PPE model mice. 70 mice total were exposed to PPE or PBS control (n = 10) at Day 1 and Day 11 to establish injury. Mice were dosed daily starting on Day 12 with nothing as a control against the placebo powder (n = 15), placebo powder (n = 15), a 1 mg dose of *Lacto* LBP (low dose, n = 15), or a 3 mg dose of *Lacto* LBP (high dose, n = 15). Half the mice were euthanized on Day 23 for a terminal sacrifice (n = 8 per group) and the other half on Day 28 for a recovery sacrifice (n = 7 per group). Safety metrics and biodistribution samples were collected throughout as described in the Table below:

| **Safety** | | **Biodistribution** | |
| --- | --- | --- | --- |
| Day 1 | Body weight, temperature | Day 2 | Urine, stool |
| Day 12 (after first dose) | Temperature (2h post), body weight | Day 12 (after first dose) | Saliva, nose swab (2h, 8h post), serum |
| Day 21 (after last dose) | Vitals | Day 21 (after last dose) | Saliva, nose swab (2h, 8h, 24h, 48h post), serum, urine, stool |
| Day 23 |  | Day 23 | Lung, brain, heart, liver, kidneys, spleen, sinuses, serum |
| Day 28 (before sac) | Vitals | Day 28 | Urine, stool (before sac)  Lung, brain, heart, liver, kidneys, spleen, sinuses, serum |

The MouseOX Plus machine (STARR Life Sciences Corp) was used to take vital signs of each mouse individually. Vital signs included: body weight, temperature, heart rate, breath rate, pulse distension, breath distension, O2 saturation, and ongoing cageside/clinical observations. Comprehensive tissue harvest was conducted to assess where, if at all, the *Lacto* LBP would distribute within the body. Bacterial gDNA was extracted from all tissues (Zymo), and PCR was run to test for the presence of the three *Lactobacillus* strains AB101, AB102, and AB103 using primers specific to each strain. Intensity of PCR products on agarose gels (1%) were quantified using ImageLab software. BAL was plated at terminal sac and recovery sac to assess if the *Lactobacillus* strains remained at the target tissue site.

*Assessment of Pulmonary Function*: At the end of hyperoxia or air exposure on PN14 (BPD model) or two days after the final LBP dose (COPD model), a subset of mice was anesthetized with isoflurane. The trachea was cannulated using a 24-gauge Angiocath and fixed with a 3-0 silk ligature. Pulmonary function was evaluated using a flexiVent apparatus (SCIREQ, Montreal, QC, Canada) that was equipped with a module 1 as described previously by Nicola et al (3). This protocol was used for all *in vivo* experiments.

*Right ventricular hypertrophy assessment*: Right ventricular (RV) hypertrophy was assessed by measuring the whole heart weight, RV free wall weight, an RV/(LV+S) ratio (Fulton index), where LV represents the left ventricle and S is the interventricular septum (4).

*Animal Harvesting*: Following PFT and/or echocardiogram, mice were euthanized and randomly assigned for lung histology (n=6), BAL (n=4-6), or lung homogenates (n=6). In order to harvest lung tissue for histology, the lungs were inflation-fixed via instillation of 0.3mL of 10% formalin through the trachea. The chest was then opened, and the lungs were harvested and placed in 10% formalin. After approximately 24 hours, the 10% formalin was exchanged for 100% isopropanol prior to being sent for processing. BAL was harvested by injecting 0.3mL of PBS via the trachea at least twice. The sample was then flash-frozen in dry ice and stored at -80°C for protein analysis. The left lower lobe of the lung was collected and processed to make whole lung homogenates (5). For COPD model mice, serum was also harvested. These tissue harvest protocols were used in all *in vivo* experiments.

*Analysis of alveolar morphometry*: After harvest, the lung tissue was sliced into 5-micrometer sections, and they were stained with hematoxylin and eosin as described in previous studies (3, 6, 7). Tissue imaging was performed with the software package MetaMorph version 6.2r4 (Universal Imaging) interfaced with a Nikon TE2000U microscope. The microscope was equipped with a QiCam Fast Cooled high-resolution CCD camera. Radial alveolar count (RAC) and mean linear intercept (MLI) were used in the evaluation of alveolar lung development (3, 7). The number of septal junctions and branches were quantified using ImageJ software as previously described (3).

*MMP-9 qPCR:* Total RNA from cells and homogenized lung tissue was extracted using TRIzol lysis reagent (15596026; Invitrogen, Waltham, MA, USA) and reverse transcribed using the SuperScript® III First-Strand Synthesis System for RT-PCR (18080-051, Invitrogen) per manufacturer’s instructions. Quantitative real-time PCR (RT-PCR) was performed using primer-probes for human MMP-9 (Hs00957562_m1, Thermo Fisher) and mouse MMP-9 per manufacturer’s instructions (Mm00442991_m1, Thermo Fisher). RT-PCR was performed on the MyiQ™ Single-Color Real-Time PCR detection System (Bio-Rad, Hercules, CA, USA) using SYBR Green PCR Master Mix per manufacturer’s instructions (4309155; Applied Biosystems, Waltham, MA, USA). RT-PCR was performed using an initial 10 min denaturation period at 95°C followed by 50 cycles of 15 s at 95°C and 1 min annealing and extension at 60°C. Expression levels of MMP-9 were normalized to 18S RNA.

*Analysis of MPO, MMP-9, CRP, IL-8 by ELISA*: Using DuoSet ELISA (R&D Systems), natural and recombinant mouse MPO, MMP-9, and CRP were measured in BAL cell pellets and serum. BAL was centrifuged at 10,000*g* for 5 minutes, pellets were resuspended in 200μl of PBS and further diluted by a factor of 10. 100μl of sample was added per well. Samples were treated with detection antibody, followed by the streptavidin-HRP method. OD was measured at 450nm in a microplate reader. Data was normalized to a standard curve.

*Analysis of cytokines*: Milliplex Mouse Cytokine Magnetic kits were custom designed (Millipore Sigma) and used to measure an array of signaling proteins. The manufacturer instructions were followed. Median fluorescent intensity (MFI) data were used for calculating cytokine/chemokine concentrations in samples, using a 5-parameter logistic method.

**Pathway Analysis**

**PCR Array**

A custom PCR array was designed (RealTime Primers; Elkins Park, PA) with 88 genes specific to human lung epithelial cells. HBEC were exposed to noxious stimuli and/or treated with the LBP for 4 hours.

1. PBS control
2. PBS + LBP
3. *Pseudomonas aeruginosa*
4. *Pseudomonas* + LBP
5. Cigarette smoke
6. Cigarette smoke + LBP
7. Smoke + *Pseudomonas*
8. Smoke + *Pseudomonas* + LBP
9. *E. coli*
10. *E. coli* + LBP

RNA was extracted using RNeasy Kit (Qiagen), and cDNA and PCR mix came from Real Time Primers.

Qiagen Ingenuity Analysis (Qiagen IPA®, Redwood City, CA) was used to generate principal coordinate analysis (PCA) and heat maps of genes with expressed altered by LBP treatment (Supplemental Figure 4).

**Supplemental Data**

**Supplemental Figure 1.**

Left) Polysaccharide (PS) isolated from *Lactobacillus* strains AB101, AB102, and AB103 increase MMP-9 levels in *E. coli* exposure model in HBEC. Right) Peptidoglycan (PD) isolated from *Lactobacillus* strains AB101, AB102, and AB103 do not affect MMP-9 levels in *E. coli* exposure model in HBEC.

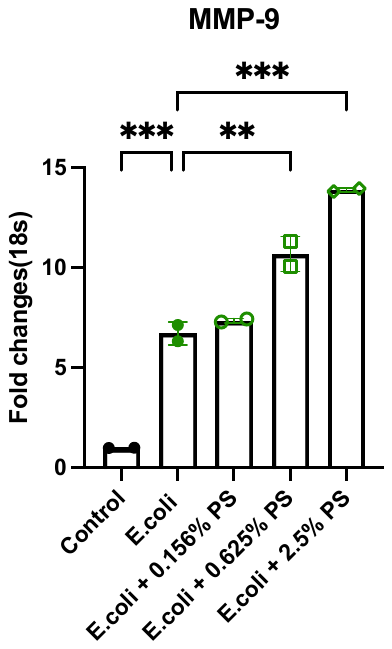

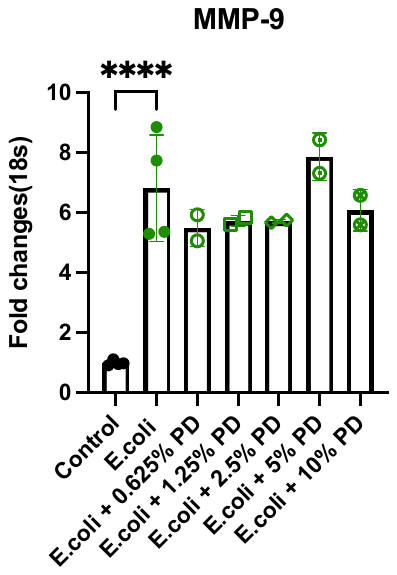

**Supplemental Figure 2.**

A) MMP-9 protein levels in serum decrease in LBP-treated (Lacto) PPE and PPE + LPS mice. B) NE protein levels in serum decrease in LBP-treated PPE and PPE + LPS mice. C) CRP protein levels in serum decrease in LBP-treated PPE + LPS mice.

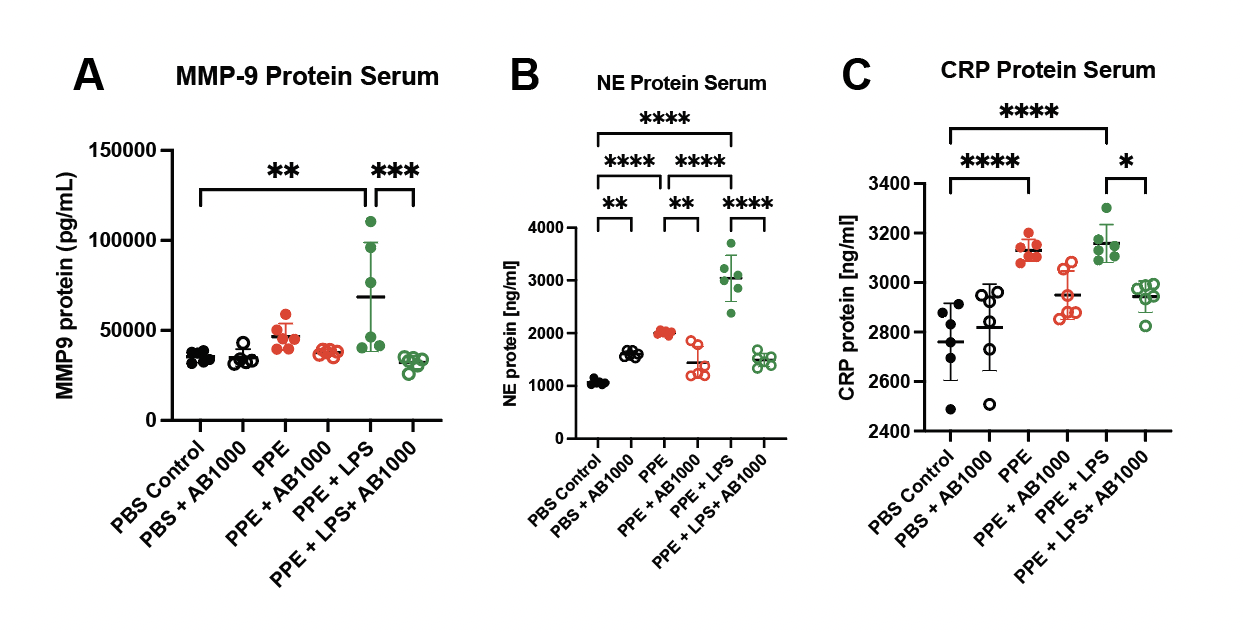

**Supplemental Figure 3.**

The *Lactobacillus* LBP performs better than steroids in reducing pro-inflammatory cytokines in the bronchoalveolar lavage fluid of mice exposed to PPE + LPS.

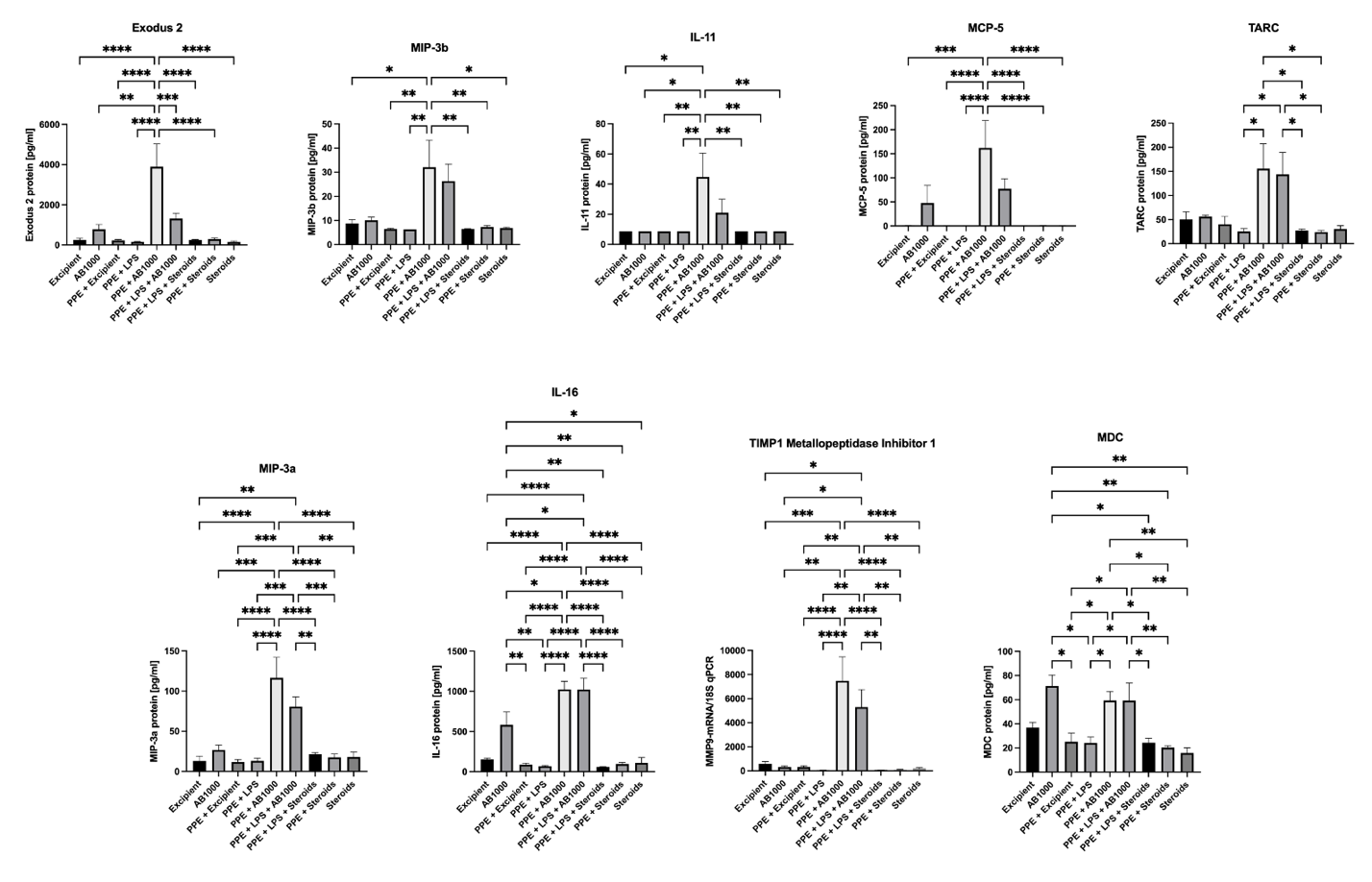

**Supplemental Figure 4.**

A) Human bronchial epithelial cells were exposed to noxious stimuli (*P. aeruginosa*, smoke, *E. coli*) and treated with LBP or a PBS control. A PCR array was conducted to identify lung epithelial-specific genes of interest with expression affected by LBP treatment.

B) Genes of interest were validated for expression in lung tissue of PPE and PPE + LPS mice (Figure 4B). MMP-9, MMP-11, HSPA4, SCN1B, CD36, CLDN18, and GRAMD2 showed a significant signal difference between injury and treatment groups.

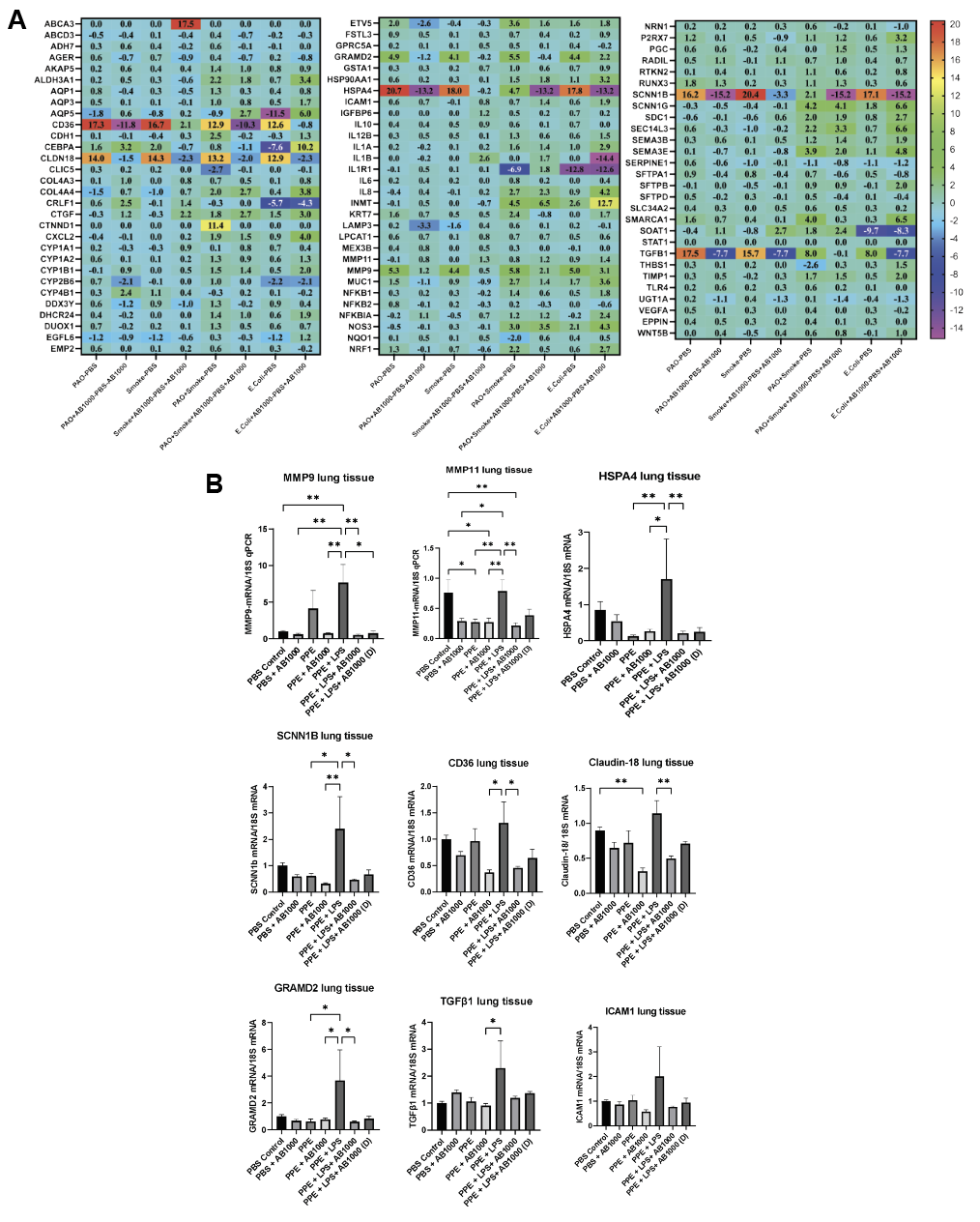

**Supplemental Figure 5.**

Batch-corrected heat map and principal coordinate analysis (PCA) comparing treatment groups of HBE cells. A) Group 1 PBS control vs. Group 2 PBS + LBP. B) Group 3 *Pseudomonas* vs. Group 4 *Pseudomonas* + LBP. C) Group 5 Smoke vs. Group 6 Smoke + LBP. D) Group 7 Smoke + *Pseudomonas* vs. Group 8 Smoke + *Pseudomonas* + LBP. E) Group 9 *E. coli* vs. Group 10 *E. coli* + LBP.

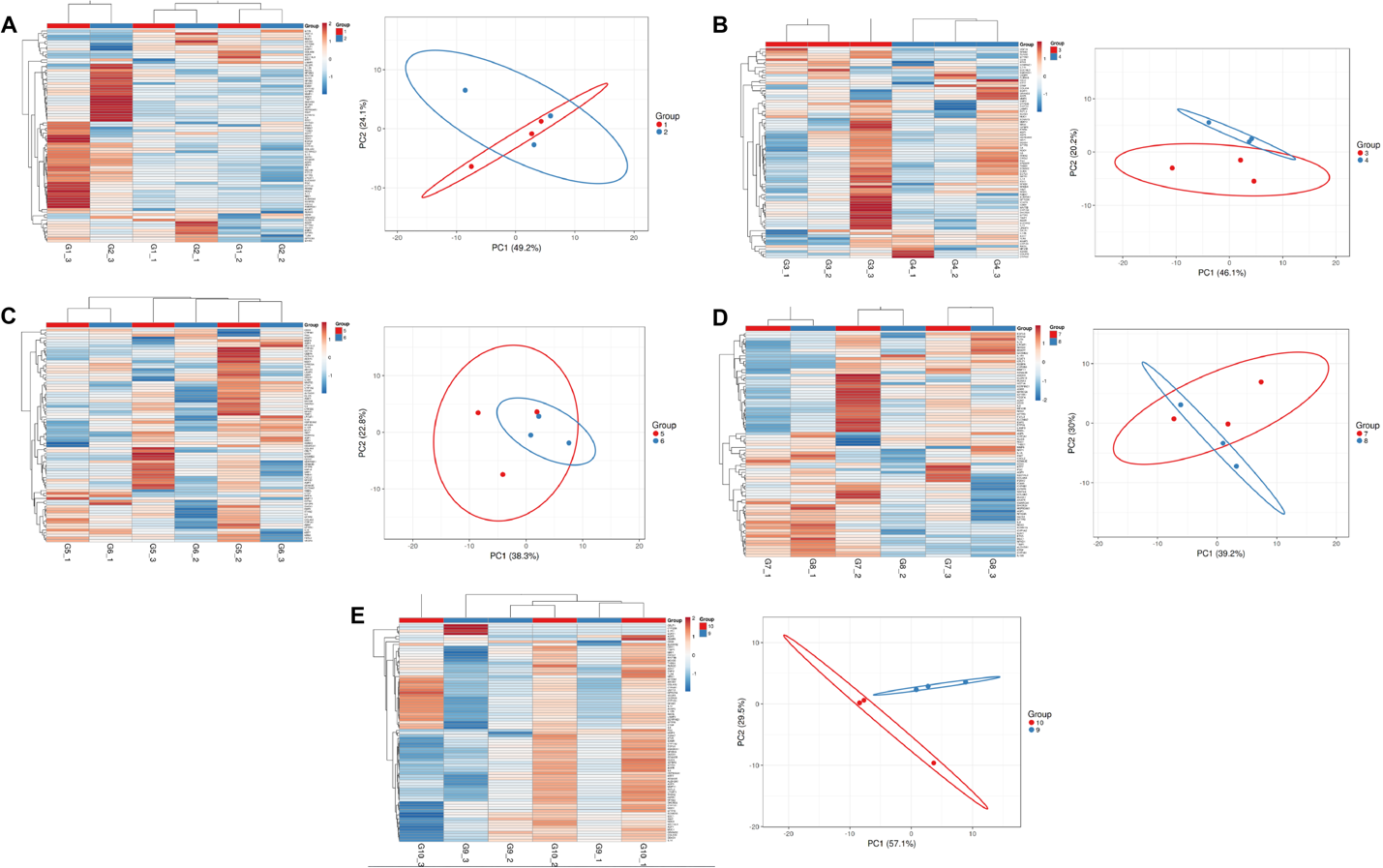

**Supplemental Table 1.** Group allocation for insufflation dose and frequency testing.

| **Group** | **Test material** | **Dose** | **Dosing Regimen** | **# Animals** | **Survival Outcome after single 5 mg dose** |
| --- | --- | --- | --- | --- | --- |
| 1 | PBS + LBP | 5 mg/2.5 mg* | Daily for 4 days | 3F | 2 died |
| 2 | PBS + LBP | 5 mg/2.5 mg* | Every other day for 4 days | 3F | 1 died |
| 3 | PPE + LBP | 5 mg/2.5 mg* | Daily for 4 days | 3F | All Survived |
| 4 | PPE + LBP | 5 mg/2.5 mg* | Every other day for 4 days | 3F | 2 died |
| 5 | LPS + LBP | 5 mg/2.5 mg* | Daily for 4 days | 3F | All Survived |
| 6 | LPS + LBP | 5 mg/2.5 mg* | Every other day for 4 days | 3F | All Survived |
| 7 | LPS | 100 µg | Day 1, Day 8 | 3F | All Survived |
| 8 | PPE | 0.25 IU | Day 1, Day 8 | 3F | All Survived |
| 9 | PBS | 50 µL | Day 1, Day 8 | 3F | All Survived |

F – female

PBS – phosphate buffered saline

PPE – porcine pancreatic elastase

LPS – lipopolysaccharide (*E. coli* origin)

*Mice were given 5 mg of LBP on Day 1. The dose was lowered to 2.5 mg on Day 2.

**Supplemental Table 2.** Per Qiagen Ingenuity Analysis, genes with decreased expression upon LBP treatment of HBEC exposed to various noxious stimuli.

|  | **Gene** | **Name** |
| --- | --- | --- |
| ***Pseudomonas* vs. *Pseudomonas* + LBP** | | |
|  | ADH7 | Alcohol dehydrogenase class 4 mu/sigma chain |
|  | CYP1A2 | Cytochrome P450 1A2 |
|  | ICAM1 | Intercellular adhesion molecule 1 |
|  | NFKB2 | Nuclear Factor Kappa B Subunit 2 |
|  | RUNX3 | RUNX Family Transcription Factor 3 |
|  | SEPARINE 1 | Serpin Family E Member 1 |
|  | SFTPA1 | Surfactant Protein A1 |
|  | WNT5B | Wnt Family Member 5B |
| **Smoke vs. Smoke + LBP** | | |
|  | AQP3 | Aquaporin 3 |
|  | CYP2B6 | Cytochrome P450 Family 2 Subfamily B Member 6 |
|  | DUOX1 | Dual Oxidase 1 |
|  | EMP2 | Epithelial Membrane Protein 2 |
|  | FSTL3 | Follistatin Like 3 |
|  | IL6 | Interleukin 6 |
|  | MMP11 | Matrix Metallopeptidase 11 |
|  | NFKB2 | Nuclear Factor Kappa B Subunit 2 |
|  | RTKN2 | Rhotekin 2 |
|  | SEMA3E | Semaphorin 3E |
|  | SFTPB | Surfactant Protein B |
|  | SLC34A2 | Solute Carrier Family 34 Member 2 |
|  | SOAT1 | Sterol O-Acyltransferase 1 |
|  | TIMP1 | TIMP Metallopeptidase Inhibitor 1 |
| **Smoke + *Pseudomonas* vs. Smoke + *Pseudomonas* + LBP** | | |
|  | CDH1 | Cadherin 1 |
|  | KRT7 | Keratin 7 |
| ***E. coli* vs. *E. coli* + LBP** | | |
|  | ABCD3 | ATP Binding Cassette Subfamily D Member 3 |
|  | ADH7 | Alcohol dehydrogenase class 4 mu/sigma chain |
|  | AKAP5 | A-Kinase Anchoring Protein 5 |
|  | CLDN18 | Claudin-18 |
|  | COL4A3 | Collagen Type IV Alpha 3 Chain |
|  | CTGF | Connective tissue growth factor |
|  | CXCL2 | C-X-C motif chemokine ligand 2 |
|  | CYP1B1 | Cytochrome P450 1B1 |
|  | CYP4B1 | Cytochrome P450 4B1 |
|  | EMP2 | Epithelial membrane protein 2 |
|  | GPRC5A | Retinoic acid-induced protein 3 |
|  | IL10 | Interleukin 10 |
|  | IL12B | Subunit beta of interleukin 12 |
|  | IL8 | Interleukin 8 |
|  | LAMP3 | Lysosome-associated membrane glycoprotein 3 |
|  | MEX3B | Mex-3 RNA Binding Family Member B |
|  | NFKB1 | Nuclear Factor Kappa B Subunit 1 |
|  | RADIL | Rap Associating With DIL Domain |
|  | RUNX3 | RUNX Family Transcription Factor 3 |
|  | SERPINE1 | Serpin Family E Member 1 |
|  | SFTPA1 | Surfactant Protein A1 |
|  | SFTPB | Surfactant Protein B |
|  | SLC34A2 | Solute Carrier Family 34 Member 2 |
|  | THBS1 | Thrombospondin 1 |
|  | TLR4 | Toll Like Receptor 4 |
|  | UGT1A | UDP Glucuronosyltransferase Fam1 Member A Complex Locus |
|  | VEGFA | Vascular Endothelial Growth Factor A |
|  | WNT5B | Wnt Family Member 5B |
